# Grow2Fold: mapping heterogeneous developmental growth to human brain folding

**DOI:** 10.64898/2026.10.08.757624

**Authors:** Jixin Hou, Akbar Solhtalab, Kun Jiang, Beikang Gu, Tiantian Li, Xianyan Chen, Taotao Wu, Kenan Song, Yang Liu, Wenzhan Song, Dajiang Zhu, Tianming Liu, Ellen Kuhl, Gang Li, Mir Jalil Razavi, Xianqiao Wang

**Affiliations:** School of ECAM, College of Engineering, University of Georgia, Athens, GA 30602, USA; Department of Mechanical Engineering, Binghamton University, Binghamton, NY 13902, USA; Department of Epidemiology and Biostatistics, College of Public Health, University of Georgia, Athens, GA 30602, USA; School of Chemical, Materials, and Biomedical Engineering, College of Engineering, University of Georgia, Athens, GA 30602, USA; School of Electrical & Computer Engineering, University of Georgia, Athens, GA 30602, USA; Department of Computer Science, University of Texas at Arlington, Arlington, TX 76019, USA; School of Computing, University of Georgia, Athens, GA 30602, USA; Department of Mechanical Engineering, Stanford University, Stanford, CA 94305, USA; Department of Radiology and Biomedical Research Imaging Center, The University of North Carolina at Chapel Hill, Chapel Hill, NC 27599, USA

**Keywords:** brain folding mechanics, heterogeneous cortical growth, prenatal brain development, developmental cortical parcellation, whole-brain biophysical modeling

## Abstract

Cortical folding emerges through complex developmental processes, yet what organizes these processes into the characteristic folding patterns of the human brain remains unclear. Here, we developed an imaging-informed whole-brain modeling framework, Grow2Fold, that integrates developmental cortical regionalization, quantitative growth characterization, and mechanics-guided simulation to investigate how cortical growth shapes brain folding patterns. Using 836 fetal and neonatal MRI sessions from 747 participants spanning 21-45 postmenstrual weeks, we identified a reproducible and hierarchical spatiotemporal organization of cortical expansion characterized by coordinated regional growth trajectories. Rapid expansion occurs earlier around peri-central territories and progressively shifts toward frontal, parietal, and temporal association cortices. We encoded these imaging-derived trajectories into region- and age-specific growth laws and incorporated them into anatomically realistic whole-brain mechanical models. The resulting simulations recapitulate major sulcal landmarks and characteristic folding development across global and regional morphological measures. Comparing regional and uniform growth, we find that differential growth *drives* cortical instability, while heterogeneous developmental growth *organizes* this instability into anatomically recognizable folding patterns. Combined with subject-specific anatomy, these population-informed growth laws reproduce individualized longitudinal folding trajectories. Together, our results identify spatiotemporally patterned developmental growth as a key spatial constraint on cortical morphogenesis and establish a quantitative bridge from developmental imaging, through mechanical instability, to anatomically organized folding.

## Introduction

Cortical folds form progressively during prenatal development, transforming the initially smooth cerebral surface into the highly convoluted architecture of the mature brain^1^. This folding process expands the cortical surface and establishes structural landmarks closely associated with brain organization and function, whereas disrupted folding has been reported across a range of neurodevelopmental conditions, including cortical malformations, autism spectrum disorder and schizophrenia^2, 3^. Although cortical fold formation involves multiple interacting biological and biochemical processes, such as progenitor proliferation, neuronal migration and molecular patterning, biomechanical mechanisms inherent to cortical growth are increasingly recognized as important contributors to its morphogenesis^4, 5^. Growth-induced mismatch in tissue deformation, in particular, can generate mechanical stress and instability that drive cortical folds^6^. Yet the cortical growth underlying these mechanical processes varies substantially across cortical regions and developmental stages, with growing genetic and transcriptomic evidence revealing region-specific developmental programs that may underlie such heterogeneity^7, 8^. A major unresolved question, therefore, is how spatiotemporally heterogeneous cortical growth interacts with tissue mechanics to organize the emergence of characteristic cortical folding patterns.

Developmental neuroimaging has enabled systematic characterization of fetal and neonatal cortical maturation. Surface-based analyses have revealed substantial regional differences in cortical expansion, indicating that cortical growth is organized into developmentally distinct territories rather than following a spatially uniform pattern^9^. This motivates the need for appropriate cortical parcellations that capture the inherent developmental organization of the cortex. Conventional parcellations are largely defined by mature anatomical landmarks or functional organization and therefore have limited applicability to the fetal cortex, particularly at early stages when majority of gyral and sulcal landmarks have not yet emerged^10, 11^. Developmental parcellation offers an alternative by grouping cortical locations according to shared growth trajectories, thereby providing a spatial representation of coordinated cortical maturation^12,13^. Reliably characterizing developmental heterogeneity, however, requires large, high-quality imaging cohorts with sufficiently dense and continuous age coverage to capture rapidly changing growth patterns across gestation^14^. Recent work has demonstrated the feasibility of this strategy by deriving developmental regionalization of cortical surface area from large-scale imaging data spanning 29 weeks postmenstrual age (PMA) to 2 years of age^13^. Nevertheless, the earlier fetal period, particularly 21-28 weeks PMA, remains poorly represented. This leaves a critical phase of cortical morphogenesis less well characterized, during which primary folds begin to emerge and the major folding architecture starts to take shape. Therefore, a more comprehensive and temporally continuous characterization of heterogeneous cortical growth spanning the broader developmental window of primary cortical folding is warranted.

Complementary to developmental neuroimaging, mechanics-based computational models provide a mechanistic framework for investigating how cortical growth drives the emergence of folding. In these models, differential growth between the expanding cortex and underlying subcortical tissue can produce incompatible deformation, generating mechanical stresses and instabilities that drive cortical folding^15-24^. Such models offer a controlled setting for testing mechanistic hypotheses that are difficult to isolate *in vivo*. However, existing approaches often rely on simplified growth prescriptions, including spatially uniform cortical expansion or predefined regional variations^15-19^, which do not fully capture the spatiotemporal heterogeneity revealed by developmental neuroimaging. In addition, many studies employ idealized or anatomically inspired brain geometries rather than MRI-derived, anatomically realistic, subject-specific representations^20-24^. This limits direct comparison with observed brain morphology and complicates quantitative evaluation against developmental imaging. These limitations highlight a persistent disconnect between developmental imaging and current biophysical models of cortical morphogenesis: imaging reveals where and when cortical growth occurs, whereas modeling provides a means to investigate how such growth shapes cortical folding. This challenge is further amplified by the limited availability of longitudinal imaging that directly captures cortical morphogenesis within individual brains. Fetal MRI remains dominated by cross-sectional observations, leaving continuous subject-specific folding trajectories difficult to characterize^25^. Consequently, despite substantial progress in both developmental imaging and computational modeling, it remains unresolved whether and how the native spatiotemporal patterning of cortical growth shapes growth-induced mechanical instabilities into characteristic cortical folding patterns. Here, we developed an imaging-informed whole-brain modeling framework, Grow2Fold, that integrates developmental neuroimaging with whole-brain mechanics to test how the native organization of cortical growth guides cortical folding morphogenesis. We derived developmental cortical parcellations and continuous growth trajectories directly from large-scale fetal and neonatal MRI spanning the major stages of cortical folding development, and used these measurements to prescribe cortical expansion in anatomically realistic computational models. We further applied the framework across longitudinal subject-specific anatomies and systematically perturbed growth, geometry, and tissue mechanics to explore its potential for generating physically grounded developmental brain trajectories when longitudinal imaging is sparse. Our analyses show that regional growth is not simply a refinement that improves morphological agreement, but a developmental constraint that influences where and how growth-induced instabilities emerge and interact. In this way, the study connects *in vivo* patterns of cortical development to their mechanical consequences and provides a mechanistic framework for investigating variability in normal and atypical cortical morphogenesis.

## Results

We developed an imaging-informed whole-brain modeling framework, Grow2Fold, to characterize spatially heterogeneous cortical development and reproduce cortical folding *in silico* (Fig. 1). Our central objective was to determine how this developmental growth heterogeneity influences the spatial organization of cortical folding. The analysis included 836 fetal and neonatal MRI sessions from 747 participants spanning 21-45 weeks of postmenstrual age (Fig. 1a; Supplementary Fig. S1; Supplementary Table S1). Across this period, cortical growth exhibited substantial regional heterogeneity. Distinct cortical regions followed different patterns and rates of surface-area expansion, providing a population-level description of the spatial organization of cortical development (Fig. 1b). We then translated these developmental patterns into region-specific growth laws for anatomically realistic whole-brain simulations (Fig. 1c). By integrating imaging-derived growth characteristics with subject-specific computational models, the framework reproduced major features of cortical morphogenesis and enabled direct quantitative comparison between simulated and anatomical brains (Fig. 1d). The framework also provides a controlled setting for testing how regional growth, geometry and mechanical conditions influence the resulting folding patterns. The following sections first define the developmental regionalization of cortical growth, then characterize regional growth dynamics, and finally evaluate their ability to reproduce cortical folding.

**Fig. 1.**
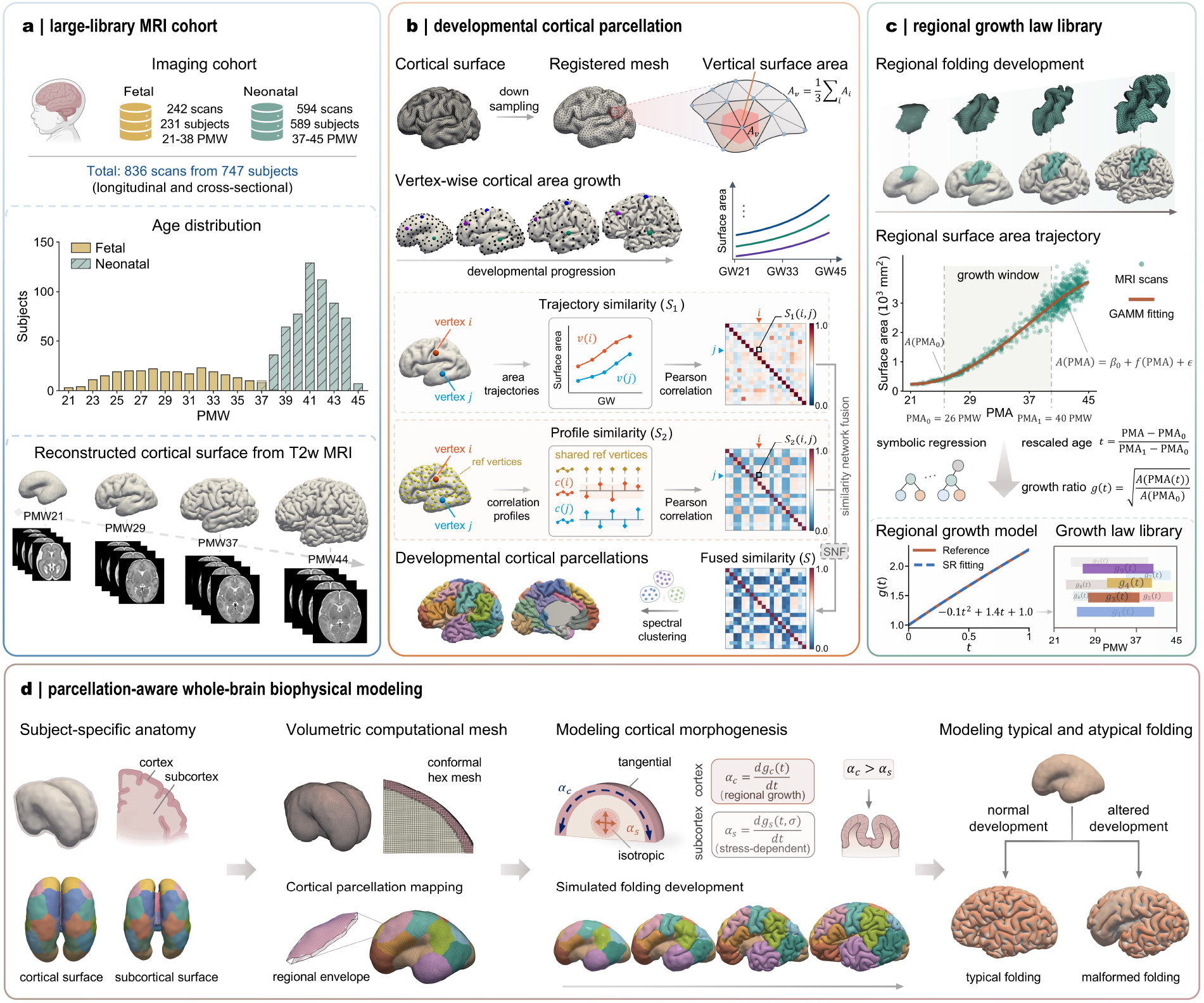
Overview of the imaging-informed whole-brain modeling platform, Grow2Fold. **a**, Large-library MRI cohort used to characterize early brain development. The dataset includes fetal and neonatal MRI scans spanning 21-45 postmenstrual weeks (PMW), with both longitudinal and cross-sectional samples. Age distributions are summarized for fetal and neonatal subjects, and representative T2-weighted MRI-derived cortical surfaces are shown across developmental stages. **b**, Developmental cortical parcellation from vertex-wise cortical surface-area growth. Cortical surfaces were down-sampled and registered to a common mesh, where local vertex-wise surface area was computed. Developmental surface-area trajectories were compared using trajectory similarity and profile similarity, and the resulting similarity networks were fused for spectral clustering to generate developmental cortical parcellations. **c**, Construction of a regional growth law library. Regional folding development and surface-area trajectories were quantified from MRI-derived cortical surfaces. Growth windows were extracted from fitted developmental trajectories, rescaled by postmenstrual age (PMA), and converted into regional growth ratios. Symbolic regression was then used to derive interpretable regional growth models, forming a growth law library across developmental windows. **d**, Parcellation-aware whole-brain biophysical modeling. Subject-specific cortical and subcortical anatomy was reconstructed and converted into conformal volumetric meshes. Cortical parcellations were mapped onto the computational mesh to assign regional growth fields. The model incorporated the regional growth laws derived in panel c, with tangential cortical growth and isotropic or stress-dependent subcortical growth, to simulate cortical morphogenesis and investigate typical and altered folding development.

### Cortical development exhibits a stable and hierarchical regional organization

To characterize the spatial heterogeneity of cortical development, we derived a developmental parcellation from vertex-wise surface-area trajectories across 21-45 weeks of postmenstrual age. Cortical vertices with similar developmental profiles were grouped by integrating trajectory-based and correlation-profile similarities, followed by spectral clustering (Fig. 1b).

Fig. 2a illustrates the progressive parcellation process as the prescribed number of clusters, *k*, increased from 2 to 25. At low model orders, the clustering first captures broad spatial divisions of the cortex. At *k* = 2, the cortex is partitioned into two large developmental territories, broadly separating the anterior/frontal cortex from the posterior cortex. As *k* increases, these coarse territories are progressively subdivided into more spatially specific regions with distinct surface-area developmental patterns (Supplementary Movie 1). Additional regions emerge primarily through subdivision of these larger territories rather than through extensive reorganization of previously identified parcels. For example, the occipital cortex separates as a distinct developmental region at *k* = 3, and its boundary remains largely preserved across subsequent clustering solutions. Similar conserved boundaries are observed in other cortical territories, including the representative examples highlighted by arrows in Fig. 2a. The persistence of these boundaries across model orders suggests that they reflect robust differences in regional developmental trajectories rather than clustering fluctuations associated with a particular choice of *k*. Although the left and right hemispheres exhibit similar early subdivision patterns, the order in which new regions emerge begins to diverge after approximately *k* = 6. Despite these intermediate differences, the two hemispheres converge toward a closely corresponding regional organization at *k* = 20. Similar nested subdivision patterns have been reported in previous growth-based fetal cortical parcellations and are consistent with a hierarchical organization of cortical development^12, 13^. The progressive subdivision and preservation of major boundaries support a stable and hierarchical organization of cortical surface-area development.

**Fig. 2.**
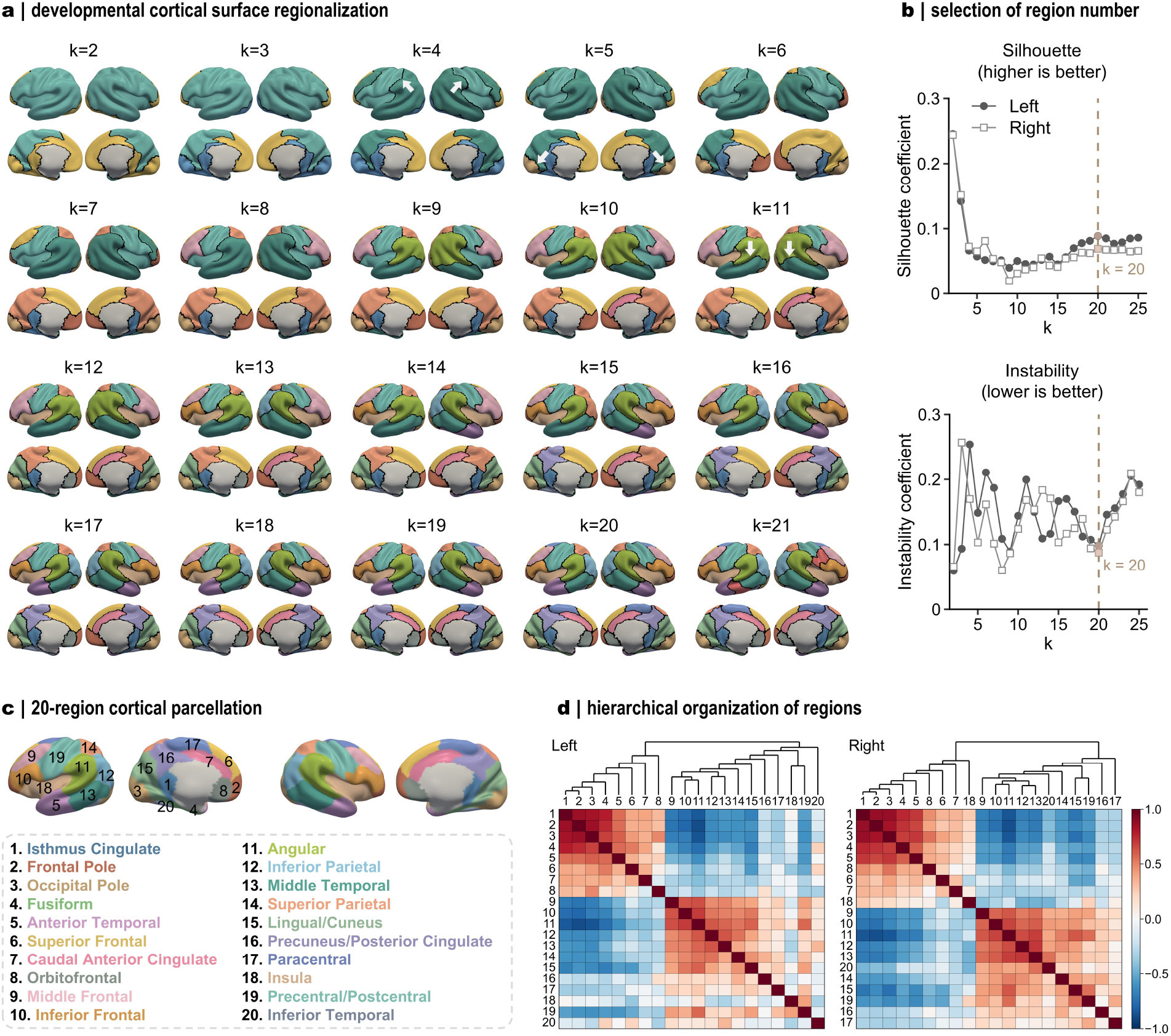
Cortical surface-area development exhibits a stable and hierarchical regional organization. **a**, Developmental cortical surface regionalization across different region numbers (clusters), k = 2-21. Cortical regions were derived from vertex-wise developmental surface-area growth patterns and visualized on lateral and medial cortical surfaces for both hemispheres. Dark lines indicate cortical regional boundaries. White arrows highlight representative boundaries that are well preserved during the spectral clustering process. **b**, Selection of the region number based on silhouette coefficient and clustering instability for the left and right hemispheres. The dashed vertical line indicates the selected model order, k = 20, which balances higher silhouette performance with lower instability. **c**, Final 20-region cortical parcellation projected onto cortical surfaces, with region indices and corresponding anatomical labels. **d**, Hierarchical organization of the 20 parcellated regions. Dendrograms and inter-regional correlation matrices show the similarity structure among regional developmental surface-area profiles for the left and right hemispheres. Color encodes the correlations between regions.

We next determined the final model order using two complementary criteria that capture clustering quality and robustness (Fig. 2b). The silhouette coefficient quantifies the separation of developmental profiles between regions, whereas split-half instability measures the sensitivity of the parcellation to variation in the imaging sample. Across candidate solutions, increasing model complexity does not consistently improve clustering quality. We selected *k* = 20 because it provided a favorable balance between a relatively high silhouette coefficient and low clustering instability in both hemispheres. Though increasing the model order to *k* = 21 introduces an additional subdivision, as shown in Fig. 2a, it does not yield a corresponding improvement in silhouette performance. Beyond *k* = 20, clustering instability also increases, indicating that further subdivisions become less reproducible across data subsets. We note that, in this study, the model order is restricted to *k* ≤ 25. Although larger model orders (*k* > 25) may potentially exhibit favorable clustering metrics, they would introduce additional anatomical subdivisions and greater regional complexity beyond the level considered here. These results support *k* = 20 as a robust parcellation solution that captures sufficient regional detail without introducing fewer stable subdivisions.

The resulting 20-region parcellation revealed spatially contiguous cortical subdivisions distributed across frontal, parietal, temporal, occipital, insular and medial cortices. We assigned anatomical descriptions to each region according to its predominant cortical location and correspondence with established anatomical landmarks (Fig. 2c). For example, region 18 delineates the insular cortex, whereas region 19 encompasses the central sulcus and adjacent precentral and postcentral cortices. Although the left and right hemispheres are parcellated independently, the resulting maps illustrate a broadly symmetric spatial organization, with homologous regions occupying comparable cortical territories on both sides. This bilateral correspondence supports the reproducibility of the developmental organization across hemispheres. We also compared the resulting parcellation with established anatomical and developmental cortical parcellations (Supplementary Table S2). The closest overall correspondence is observed with the Huang developmental surface-area parcellation^13^, followed by the Wang cortical-thickness parcellation^26^ and the FreeSurfer adult anatomical parcellation^27^. Quantitative comparisons based on the adjusted Rand index (ARI), normalized mutual information (NMI), and Dice overlap are provided in the Supplementary Materials. The present parcellation retains distinct regional boundaries and spatial subdivisions, reflecting differences in cohort composition, postmenstrual age range, and regionalization strategy.

We further examined the relationships among the identified regions using correlations between their developmental surface-area profiles (Fig. 2d). The inter-regional correlation matrices revealed groups of regions with strongly coordinated developmental trajectories, while the corresponding dendrograms organized these regions into progressively nested branches. In both hemispheres, the regions were organized into two major developmental clusters, although the detailed composition and branching structure differed slightly between hemispheres. Within each cluster, regions with more similar trajectories formed progressively finer subgroups. This hierarchical organization indicates that cortical surface-area development is regionally coordinated rather than spatially uniform. Together, these results establish a robust developmental parcellation that captures coordinated regional cortical growth and provides the spatial basis for the subsequent regional growth characterization and computational modeling.

### Regional cortical expansion exhibits distinct spatiotemporal developmental patterns

Using the developmental parcellation defined above, we next characterized regional cortical surface-area growth from 21 to 45 weeks PMA (Fig. 3). For each region, surface area increases continuously with postmenstrual age, but the magnitude and temporal profile of expansion vary substantially across the cortex (Fig. 3a). To capture the regional growth trends across development, we fitted surface-area trajectories using generalized additive mixed models (GAMMs). This approach flexibly captures nonlinear age-related changes without imposing a predefined growth function and is well suited to developmental datasets with uneven age sampling. The fitted trajectories show relatively steady, near-linear expansion in most regions from approximately 21 to 28 weeks PMA, followed by more rapid growth during the late second and third trimesters. The degree and timing of this acceleration differ among regions, indicating pronounced spatial heterogeneity in cortical surface-area development. Similar developmental patterns are also observed in the right hemisphere (Supplementary Fig. S2).

**Fig. 3.**
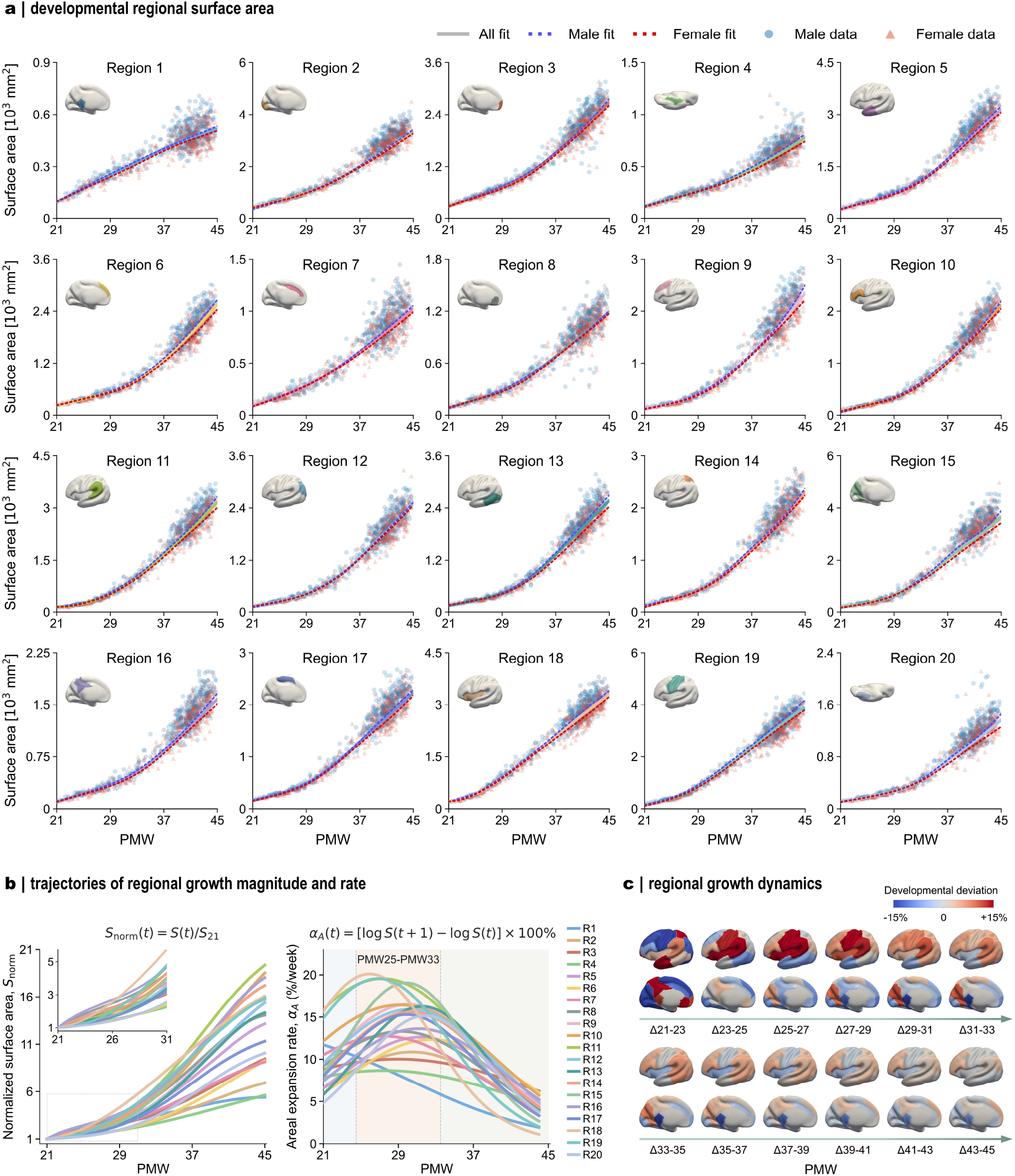
Regional cortical expansion exhibits distinct spatiotemporal developmental patterns. **a**, Regional cortical surface-area trajectories from 21 to 45 postmenstrual weeks (PMW) across 20 cortical regions. Blue circles and red triangles denote male and female data, respectively; solid grey curves denote fits across all subjects, and dashed blue and red curves denote sex-specific fits. Insets show the anatomical locations of the corresponding regions. **b**, Trajectories of regional growth magnitude and rate derived from the fitted surface-area curves. Left, surface-area trajectories normalized to 21-week PMA show accumulated regional expansion, whereas right, areal expansion rates capture the temporal dynamics of regional growth speed. **c**, Spatial maps of regional growth dynamics across developmental windows. Each cortical region is assigned the deviation of its interval-specific expansion percentage from the mean regional expansion within the same window. Red and blue indicate above-average and below-average growth behaviors, respectively.

We additionally assessed hemispheric and sex-related differences in regional cortical surface-area development. Although the developmental parcellation presents broadly similar spatial organization across hemispheres, homologous regions exhibit region-specific differences in cortical surface area (Supplementary Table S3). We quantified these differences using the surface-area asymmetry index (SAI), with positive and negative values indicating leftward and rightward asymmetry, respectively. Both the magnitude and direction of asymmetry vary across regions, indicating spatially heterogeneous left-right differences rather than a uniform hemispheric bias. For sex-related differences, fitted trajectories generally show larger absolute regional surface areas in males than in females across development (Fig. 3a). We further quantified the onset of these differences and found that significant effects emerged at region-specific ages, beginning at approximately 30-week PMA in several regions (Supplementary Table S4). To isolate regional effects from differences in overall cortical size, we repeated the analysis with total cortical surface area included as a covariate. After adjustment, most sex differences are no longer significant, and the remaining effects are limited to a small subset of regions and generally appear later in development. These findings suggest that much of the observed sex difference in regional surface area is associated with overall cortical size rather than widespread divergence in regional developmental trajectories.

To compare developmental pace across regions, we normalized each fitted surface-area trajectory to its value at 21-week PMA and derived the corresponding areal expansion rate (Fig. 3b). The normalized trajectories diverge progressively with age, reflecting substantial differences in accumulated regional expansion. The expansion-rate curves further reveal a concentrated period of rapid cortical growth between approximately 25- and 33-week PMA, although the timing and magnitude of the peak varied among regions. This developmental interval coincides with pronounced changes in cortical morphology, during which primary sulci become established and cortical folding increases rapidly. To further characterize regional growth heterogeneity, we mapped relative cortical expansion, defined as each region’s deviation from the mean cortical expansion within the same interval, across developmental windows (Fig. 3c). Before approximately 29-week PMA, above-average expansion is concentrated around the central and peri-central cortices, including territories surrounding the developing central sulcus. These regional differences become less pronounced during the subsequent interval before the dominant growth pattern shifted toward frontal, parietal and temporal cortices at later ages. This spatial progression is consistent with previous longitudinal observations showing that rapid cortical expansion shifts from primary sensorimotor regions toward parietal, temporal and frontal association cortices over development^28^. Together, these results demonstrate that cortical expansion is both regionally heterogeneous and temporally dynamic, providing the quantitative basis for constructing region- and age-specific growth laws for the subsequent computational models.

### Regional growth-guided simulations reproduce anatomically organized cortical folding

To translate the population-level developmental trajectories into growth controls for computational modeling, we constructed a regional growth-law library spanning cortical regions and developmental windows. For any selected interval, the fitted regional surface-area trajectory was converted to a tangential growth ratio and normalized by the surface area at the starting age. Postmenstrual age within each selected window was rescaled to a dimensionless simulation time from 0 to 1. We then used symbolic regression to convert the normalized growth trajectories into explicit polynomial functions. These analytical expressions are readily differentiable and can be directly implemented in the computational model. Because growth laws can be generated from any interval of the continuous GAMM trajectories, the library is not restricted to a fixed developmental window. This flexibility is particularly useful for subject-specific modeling, where growth laws can be constructed according to the ages of the available initial and follow-up MRI scans. As an example, Supplementary Fig. S3 shows the regional growth trajectories and symbolic-regression fits for the 22-42 weeks PMA window. As seen, the analytical functions closely match the imaging-derived reference trajectories across all 20 cortical regions.

With the imaging-derived regional growth laws established, we are able to incorporate them into finite-element simulations of cortical morphogenesis (Fig. 4). During modeling, we prescribed region-specific tangential cortical growth within a differential-growth framework to drive the emergence of cortical folding. Fig. 4a shows representative cortical surfaces from 21-to 45-week PMA. The anatomical brains represent cross-sectional observations from different individuals, whereas each simulated sequence follows the longitudinal evolution of the same computational brain from its initial configuration. As growth progresses, the initially smooth simulated cortex develops increasingly complex gyral and sulcal morphology, recapitulating the overall developmental progression observed in the anatomical cohort (Supplementary Movie 2). This longitudinal representation enables continuous tracking of intermediate folding states, which remain difficult to observe directly *in vivo* because fetal neuroimaging datasets are predominantly cross-sectional and repeated longitudinal acquisitions remain limited^1^. We further evaluated whether the simulations reproduced characteristic folding landmarks (Fig. 4b). Eight major sulci were extracted for comparison, including the central, precentral, postcentral, superior frontal, inferior frontal, lateral, intraparietal and superior temporal sulci. Across three representative simulations, these landmarks emerge in anatomically matched cortical territories and show comparable spatial extent and orientation to those observed in the reference brain. Major sulci are informative anatomical landmarks because primary folding patterns are relatively conserved across individuals and closely reflect the structural and functional organization of the cortex^29^.

**Fig. 4.**
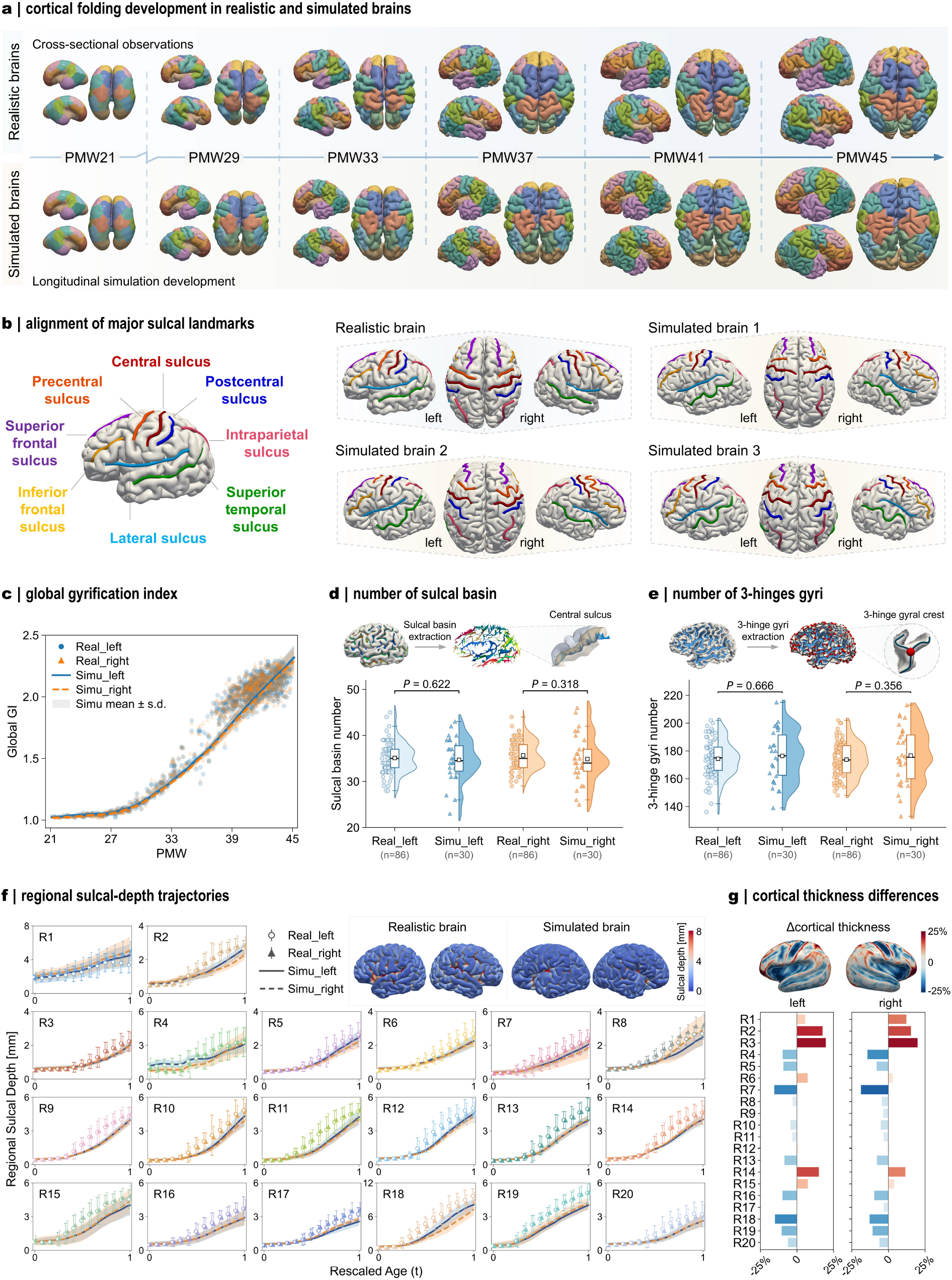
Regional growth-guided whole-brain simulations reproduce key features of prenatal cortical folding. **a**, Folding development in representative realistic and simulated brains from 21 to 45 postmenstrual week (PMW). At each postmenstrual age, lateral views of the left and right hemispheres and a whole-brain view are shown. Realistic brains are cross-sectional observations, whereas simulated brains show the longitudinal evolution of the same computational model. Cortical colors denote the 20-region developmental parcellation. **b**, Alignment of major sulcal landmarks between a representative realistic brain and three simulated brains. The central, precentral, postcentral, superior frontal, inferior frontal, lateral, intraparietal and superior temporal sulci are highlighted. **c**, Developmental trajectories of the global gyrification index (GI) in the left and right hemispheres of realistic and simulated brains. **d**, Comparison of the number of sulcal basins between realistic and simulated hemispheres. The schematic illustrates the extraction of sulcal basins and a representative central sulcus. **e**, Comparison of the number of 3-hinge gyri between realistic and simulated hemispheres. The schematic illustrates the extraction of 3-hinge gyral configurations and a representative 3-hinge gyral crest. In **d** and **e**, half-violin plots show the distributions, box plots indicate the median and interquartile range, and individual points denote brain hemispheres; sample sizes are shown below each group. Reported *P* values correspond to comparisons between realistic and simulated brains in terms of different hemispheres. Statistical significance was assessed using two-sided permutation tests of the difference in arithmetic means with 100,000 permutations. **f**, Regional sulcal-depth trajectories for the 20 cortical regions. Representative spatial maps compare sulcal-depth distributions in realistic and simulated brains. Age was rescaled to the interval 0-1. **g**, Differences in cortical thickness between realistic and simulated brains, shown as surface maps and region-wise percentage differences for the left and right hemispheres. Percentage difference was calculated as (T_simu_-T_real_)/T_real_× 100%, such that positive values indicate greater cortical thickness in the simulated brains.

We further quantified the agreement between simulated and anatomical folding using complementary global and regional measures (Fig. 4c-g). We first assessed overall folding complexity using the global gyrification index (GI), defined as the ratio of cortical surface area to the area of its convex hull. GI increased progressively with postmenstrual age in both anatomical and simulated brains, and the simulated trajectories closely tracked the developmental range observed in the imaging data for both hemispheres. Beyond the global measure, we also evaluated key folding features, including the major sulcal basins and 3-hinge gyri (Fig. 4d, e). Major sulcal basins are identified as connected cortical regions with sufficiently negative mean curvature after removing small isolated components^30^, whereas 3-hinge gyri are defined as junctions where three gyral crest branches converge within the principal gyral crest network^31^. These structures, particularly 3-hinge gyri, have been proposed as informative cortical landmarks because of their conserved structural organization and distinctive connectivity patterns^32^, and have also been investigated as potential biomarkers of cortical organization and neurological disorders^33, 34^. Both the sulcal-basin and 3-hinge gyral counts show comparable distributions between anatomical and simulated hemispheres, with no significant differences in group means (Fig. 4d, e).

To provide a more spatially resolved assessment of folding development, we quantified regional sulcal depth and cortical thickness (Fig. 4f, g). Across the 20 cortical regions, simulated sulcal-depth trajectories closely match the developmental trends and spatial patterns observed in the anatomical brains. Moreover, the sulcal-depth magnitudes are also reasonably captured through most developmental stages, with modestly lower values in several simulated regions near the final stage (Fig. 4f). For regional cortical thickness, the percentage difference between simulated and anatomical brains remains within approximately 25% across both hemispheres (Fig. 4g). Relatively higher thickness predictions are observed in R2 (frontal pole) and R3 (occipital pole), whereas lower predictions occur in regions such as R7 (caudal anterior cingulate) and R18 (insula). These regional discrepancies may arise from local contact effects during mechanical folding, which complicate the reproduction of cortical thickness at specific locations.

Together, these qualitative and quantitative comparisons support that our computational framework is capable of reproducing anatomically faithful cortical folds and key features of cortical morphogenesis with notable fidelity across multiple anatomical and morphological measures.

### Growth, geometry, and tissue mechanics shape individual and atypical cortical folding

In this section, we demonstrate several representative capabilities of the modeling framework, including subject-specific longitudinal simulation, flexible generation of cortical folding phenotypes, and mechanistic investigation of atypical cortical development (Fig. 5).

**Fig. 5.**
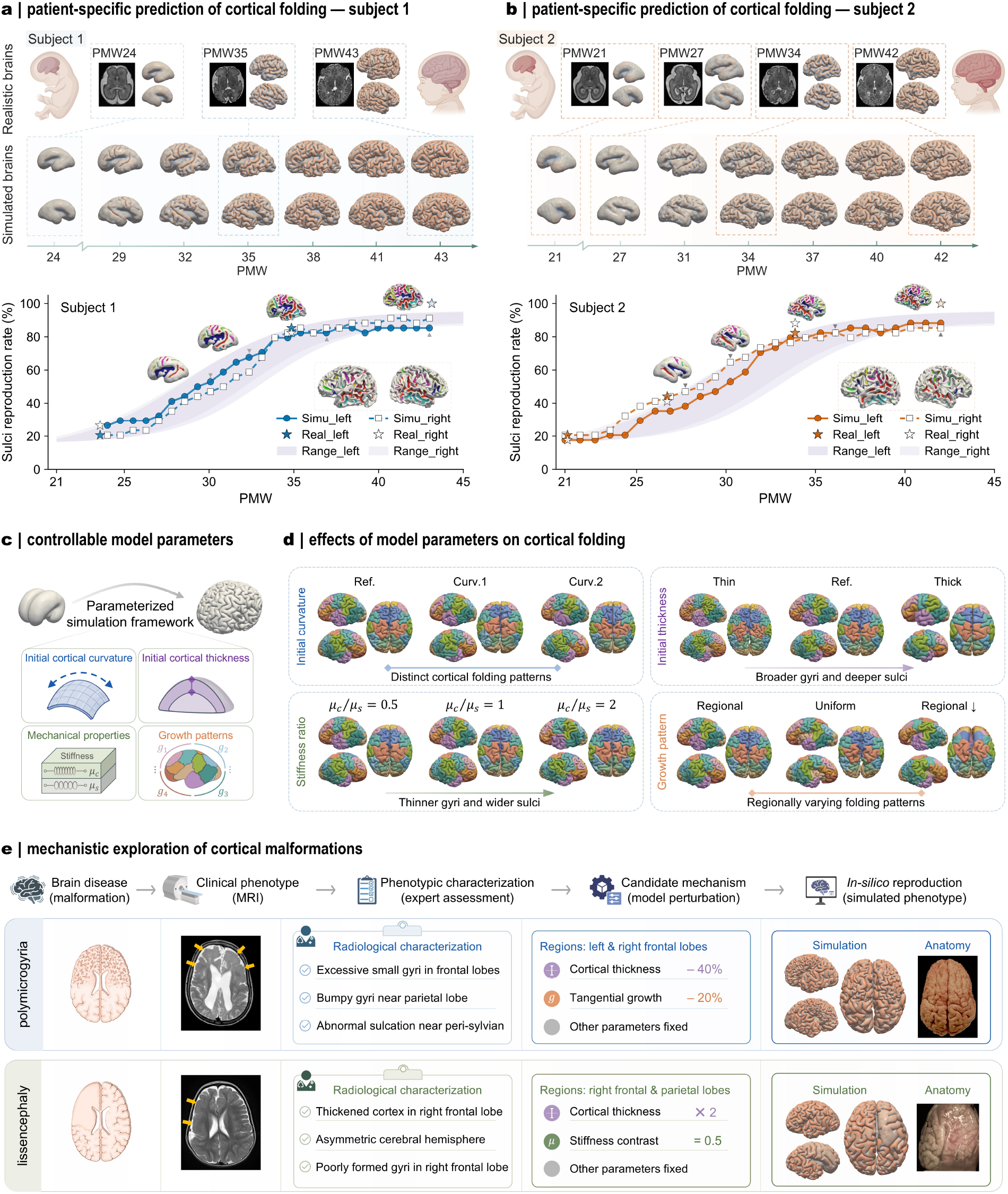
The modeling framework demonstrates capabilities for longitudinal simulation, parametric exploration and mechanistic investigation of cortical malformations. **a, b**, Patient-specific longitudinal prediction of cortical folding for subjects CC00990XX18 (**a**) and CC01019XX13 (**b**) from the developing Human Connectome Project (dHCP) dataset. Subject-specific MRI and cortical surfaces at selected postmenstrual weeks (PMW) are shown above the corresponding longitudinal simulations. Sulcal reproduction rates are quantified separately for the left and right hemispheres as (N_simu_/N_real_ *100%), where N_real_ is the number of primary sulci defined from the Destrieux parcellation in the mature reference brain. The final longitudinal anatomical brain (bottom right) is used as the reference for calculating the reproduction rate. Sulcal distributions of simulated brains at four representative time points (gray triangles) are shown above the curves. Shaded bands indicate the range obtained from ten simulated longitudinal cases. **c**, Controllable parameters in the whole-brain simulation framework, including initial cortical curvature, cortical thickness, mechanical properties and regional growth patterns. Mechanical properties can include stiffness, bulk modulus, and other constitutive quantities; shear stiffness is shown as a representative example. **d**, Representative changes in cortical folding induced by varying individual model parameters. Curv.1 and Curv.2 denote initial geometries generated by applying spatially smoothed random out-of-plane perturbations to the reference surface, with amplitudes of 0.4 and 0.8 mm, respectively. Thin and Thick denote cortical surfaces shifted inward and outward by 0.5 mm relative to the reference thickness. The stiffness-ratio cases vary the cortex-to-subcortex shear-modulus ratio. Regional growth assigns region-specific growth models from the growth library, Uniform growth assigns identical linear growth (g(t) =1.82*t) across all regions, and Regional ↓ denotes selective reduction of growth in specified regions while preserving the reference regional pattern elsewhere. Left-hemisphere, right-hemisphere and whole-brain views are shown for each condition. **e**, Mechanistic exploration of cortical malformations using polymicrogyria and lissencephaly as representative examples. Clinical MRI phenotypes are summarized through expert radiological characterization and translated into targeted perturbations of model parameters while other parameters are held fixed. The resulting *in silico* phenotypes are compared with corresponding anatomical observations, illustrating the potential of the framework for mechanism-informed investigation of abnormal cortical development. MRI images: polymicrogyria, adapted from ref.^37^ under the CC BY 4.0 license; lissencephaly: courtesy of Ammar Haouimi, Radiopaedia.org (ID: 67632). Anatomy images: polymicrogyria, adapted from ref.^38^ under the CC BY 4.0 license; lissencephaly, adapted from ref.^39^ under the CC BY 4.0 license. Illustrative icons were created using BioRender. Ref., reference configuration; Curv., curvature perturbation; (µ_c_) and µ_s_), cortical and subcortical shear moduli, respectively.

Beyond reproducing population-level cortical folding, the framework can be adapted to subject-specific longitudinal modeling when repeated MRI scans are available (Fig. 5a, b). For each subject, the initial MRI provides the anatomical geometry for constructing the computational model, while the interval between the initial and follow-up scans defines the developmental window used to retrieve regional growth laws from the growth library. We demonstrate this capability in two representative dHCP subjects with three and four longitudinal examinations spanning 24-43 weeks PMA and 21-42 weeks PMA, respectively. At each available imaging age, the corresponding MRI and reconstructed cortical morphology are shown together with the longitudinal simulation, enabling direct comparison of folding development over time. The simulated brains progressively develop major gyral and sulcal structures that are spatially consistent with those observed in the longitudinal anatomical scans (Supplementary Movies 3 and 4). The overall folding sequence is also preserved across developmental stages, although local discrepancies remain. For example, at intermediate ages such as 34-35 weeks PMA, the simulations exhibit more complex folding patterns than the anatomical brains. Additionally, the increase in folding complexity from the intermediate to final stages is less pronounced in the simulations than in the corresponding longitudinal scans. To quantify subject-specific folding reproduction, we focused on primary sulci, defined by the Destrieux atlas^27^, and assessed their reproduction based on the spatial overlap between detected sulcal basins and the corresponding atlas regions. As shown, the reproduction rate progressively increases during simulation. Moreover, the simulated trajectories show close agreement with the reproduction rates measured at the available longitudinal MRI time points and reach values above 80% at the mature stage in both representative subjects. At the final stage, residual differences remain because some sulci are only partially reproduced or are spatially misaligned with the corresponding anatomical regions. These results demonstrate the framework’s ability to generate individualized longitudinal folding trajectories from subject-specific anatomy and developmental growth information.

We next show that the framework can flexibly generate diverse cortical folding phenotypes, highlighting its controllability and potential scalability for future synthetic brain generation (Fig. 5c, d). The model allows independent modification of several key parameters, including the initial cortical geometry, cortical thickness, tissue mechanical properties and regional growth patterns (Fig. 5c). As shown in Fig. 5d, altering the initial surface curvature changes both the locations and configurations of the resulting folds, highlighting the sensitivity of cortical morphogenesis to the initial geometry. Variations in cortical thickness also produce distinct folding phenotypes: a thicker cortex generates broader and more prominent gyri with deeper sulci, whereas a thinner cortex produces smaller and shallower folds. Tissue mechanical properties can likewise be varied, as illustrated here by changes in the cortex-to-subcortex shear stiffness ratio. Increasing the stiffness contrast produces sharper gyri and wider intervening sulci. Notably, the cortical and subcortical tissues in the reference model have comparable stiffnesses, consistent with experimental measurements^35^, and this condition yields a simulated folding pattern that more closely resembles the anatomical brain, particularly in the organization of major gyral crests. These trends are consistent with mechanical buckling principles for layered core-shell systems, in which cortical thickness and stiffness contrast regulate the characteristic wavelength and morphology of surface instabilities^36^.

The framework also enables direct manipulation of cortical growth patterns. Compared with region-specific growth, imposing uniform growth across the cortex produces a distinct folding configuration. The difference is particularly evident around the insular and adjacent cortical territories. Under uniform growth, the surrounding cortical regions remain more widely separated and do not achieve the close apposition observed in the anatomical brain, resulting in an abnormally open morphology around the insula (Fig. 5d; Supplementary Fig. S4a). In contrast, regional growth produces a more compact folding configuration in this area. Consistent with this qualitative improvement, the regional-growth model also shows closer quantitative agreement with the anatomical data in global gyrification trajectories (Supplementary Fig. S4b), regional local gyrification (Supplementary Fig. S4c), and regional sulcal-depth development (Supplementary Fig. S4d), with reduced deviations relative to the uniform-growth model across many cortical regions. These results indicate that cortical growth does more than drive folding instability: its spatial heterogeneity contributes to organizing the resulting folds into anatomically recognizable configurations and improves the reproduction of both global and regional characteristics of cortical folding^21, 40^. Additionally, growth magnitude can also be selectively increased or reduced within individual regions, providing a controlled means of examining how localized developmental perturbations reshape cortical morphology. Together, these parameter studies show how growth, geometry and tissue mechanics jointly regulate cortical folding and highlight the distinct role of regional growth heterogeneity in its spatial organization.

Finally, we demonstrate that the framework can be used to investigate cortical malformations from a mechanics-informed perspective (Fig. 5e). Cortical malformations arise from abnormal developmental processes that disrupt the formation and organization of cortical folds^41^. In Fig. 5e, we use polymicrogyria and lissencephaly as two representative examples to demonstrate how such pathological morphologies can be examined within the modeling framework. Based on their characteristic radiological features, we translated the observed cortical abnormalities into corresponding geometric, mechanical and growth-related model perturbations. For the polymicrogyria example, the characteristic pattern of numerous small, closely spaced folds was reproduced by reducing cortical thickness by 40% and regional growth by 20% within the affected territory. By contrast, the smoother cortical morphology associated with lissencephaly was reproduced by doubling regional cortical thickness and reducing the local cortex-to-subcortex stiffness contrast by 50%. These perturbation magnitudes were selected to qualitatively reproduce the characteristic radiological features of the corresponding clinical cases rather than to represent quantitatively established disease-specific parameters. Because subject-specific MRI geometries were unavailable for these clinical examples, the comparisons illustrate phenotype-level correspondence rather than patient-specific reconstruction. Nevertheless, these demonstrations suggest that the framework could provide a controlled platform for evaluating candidate mechanical contributors to abnormal cortical development and, when longitudinal patient imaging is available, for exploring possible follow-up morphological trajectories.

## Discussion

Cortical folding is a complex morphogenetic process governed by coordinated biological, biochemical and mechanical mechanisms across multiple spatial and temporal scales^4^. Processes including progenitor proliferation, neuronal migration, molecular patterning and tissue growth collectively shape the developing cortex^42^. From a mechanical perspective, large-scale cortical folding can be understood as arising from the interaction between spatially patterned tissue growth and the mechanical constraints of the developing brain^43, 44^. Resolving this interaction requires connecting developmental growth measured *in vivo* with its mechanical consequences during morphogenesis. Yet developmental neuroimaging and biomechanical modeling have largely remained separate: imaging reveals where and when the cortex grows, whereas computational models explain how differential growth can generate folds. Here, by integrating large-scale developmental neuroimaging with anatomically realistic whole-brain simulation, we show that spatially heterogeneous developmental growth provides regional biases that help organize growth-induced mechanical instability into anatomically recognizable folding patterns. We identify regions based on shared developmental growth patterns and use their population-level trajectories to prescribe region-specific growth in individual brain geometries.

The developmental parcellation and regional growth trajectories support the view that cortical maturation is organized through spatially structured and temporally evolving developmental programs. The progressive subdivision of broad cortical territories, together with the persistence of major regional boundaries across clustering resolutions, suggests that cortical development is hierarchically organized rather than partitioned into independent local units. This organization is also dynamic across gestation: relatively rapid expansion is concentrated earlier around the central and peri-central cortices and later shifts toward frontal, parietal and temporal association regions. Such spatiotemporal gradients reflect underlying biological heterogeneity rather than purely geometric differences. Consistent with this interpretation, prenatal cortical development is characterized by region- and layer-specific transcriptional programs, spatial variation in neural progenitor populations, and temporally regulated cellular differentiation^45^. Spatial transcriptomic studies further show that regional transcriptional specialization and areal boundaries are already evident during fetal development^8^, while outer radial glia and related progenitor populations are strongly implicated in tangential cortical expansion^46^. These molecular and cellular processes provide a plausible biological basis for the region-specific tangential growth patterns observed by neuroimaging. Moreover, the developmental shift from earlier peri-central expansion toward later association cortices is also broadly compatible with the known progression from earlier maturation of primary sensorimotor territories toward later-developing association regions^28^. Thus, the present parcellation should not be interpreted simply as a geometric division of the cortical surface, but as a mesoscale representation of coordinated developmental programs that evolve across gestation.

A central finding of this study is that differential growth *drives* cortical instability, while heterogeneous developmental growth helps *organize* this instability into anatomically recognizable folding patterns. Differential cortical expansion can generate folding, but uniform growth provides limited spatial information about where anatomically recognizable folds should emerge^18, 20^. By incorporating imaging-derived regional growth, the model introduces spatially varying growth mismatch across the cortex, thereby influencing the locations and interactions of emerging folds. This effect is particularly evident in anatomically complex regions such as the insular and peri-Sylvian cortex, where uniform growth produces poorly formed folding configurations. Regional growth should therefore not be viewed merely as a refinement that improves morphological agreement; rather, it provides developmentally meaningful spatial constraints that help transform generic mechanical instability into folding patterns resembling those of the developing human brain. Cortical folding is shaped by the interplay of growth, geometry and tissue mechanics rather than by growth alone^47^. Initial cortical geometry, cortical thickness and material properties influence how growth-induced instabilities develop. Cortical folding may therefore be understood as the outcome of several interacting constraints: developmental growth provides the driving force, its spatial heterogeneity provides regional organization, early anatomy defines the geometric landscape in which instability develops, and tissue mechanics regulates the characteristic scale and expression of folding. This framework offers a useful explanation for why major sulci can remain relatively conserved across individuals while smaller secondary and tertiary folds show much greater variability.

The continuous representation of regional growth makes the framework well suited for subject-specific developmental modeling. Because growth laws can be derived for different developmental windows, simulations can be initialized from MRI acquired at different ages and evolved over the corresponding intervals. In the longitudinal examples, subject-specific anatomy combined with population-derived growth information reproduced the progressive emergence of major sulcal structures across repeated scans. At present, this approach is more appropriately described as population-informed subject-specific modeling, because the anatomy is individualized whereas the growth prior remains population based. Nevertheless, it provides a practical foundation for personalized developmental prediction. With denser longitudinal imaging, subject-specific growth measurements could progressively replace population priors, enabling increasingly individualized forecasts of cortical morphogenesis.

Beyond reproducing normal development, the framework offers broader opportunities for developmental modeling and translational investigation. Subject-specific simulations can interpolate developmental trajectories between sparse MRI observations, which is particularly valuable when repeated fetal imaging is difficult to obtain. Additionally, systematic tuning of geometry, growth and material parameters enables the generation of diverse yet physically plausible cortical morphologies. Such synthetic brains retain known developmental inputs and mechanistic provenance, making them potentially useful for training and validating AI models of cortical morphogenesis and developmental prediction. Recent studies have explored individualized cortical development prediction through transfer learning^48^, disentangled representation learning^49^, surface-based neural modeling^50^, and implicit neural modeling^51^. However, these approaches are often limited by the scarcity and irregular sampling of longitudinal infant imaging, resulting in insufficient paired observations for learning temporally consistent developmental trajectories. Mechanics-based generation of high-fidelity developmental brain morphologies could therefore augment sparse longitudinal imaging and provide richer training data for AI models of cortical morphogenesis. The framework further provides a mechanical perspective on cortical malformations. Polymicrogyria-like and lissencephaly-like morphologies can arise from distinct combinations of altered cortical thickness, regional growth and mechanical contrast, illustrating how different developmental perturbations may converge on characteristic abnormal folding phenotypes. With patient-specific longitudinal data, such models could ultimately complement imaging-based assessment by supporting early risk evaluation, investigation of plausible disease progression, and prediction of follow-up morphology.

Several aspects of the framework can be further refined. The current 20-region representation captures robust mesoscale growth organization but inevitably averages over finer spatial heterogeneity. Future implementations could incorporate continuous or vertex-wise growth fields while preserving the broader developmental organization identified here. In addition, cortical surface-area expansion is used as the primary imaging-derived measure of tangential growth, although the underlying developmental process also reflects neuronal proliferation and migration, cortical-layer formation, axonal organization and extracellular remodeling. Integrating complementary imaging, histological or molecular measurements could therefore provide a richer description of regional growth. The mechanical model could also be extended to incorporate greater spatial and temporal variation in tissue properties, detailed white-matter architecture, and transient fetal structures, which may further improve the representation of local folding mechanics.

Together, this work establishes a quantitative link between heterogeneous developmental growth and the mechanical organization of prenatal cortical folding. Our results suggest that differential growth *drives* the instability underlying cortical folding, while spatial heterogeneity in developmental growth *organizes* this instability across the cortex. By integrating developmental regionalization, regional growth and subject-specific anatomy, the framework provides a basis for population-informed subject-specific prediction, physically grounded synthetic brain morphologies, and mechanics-informed investigation of atypical cortical development.

## Methods

Our pipeline links developmental neuroimaging with growth characterization and physics-based modeling of cortical folding. It integrates fetal and neonatal MRI processing, data-driven cortical parcellation, regional growth-law construction, subject-specific finite-element simulation, and quantitative evaluation of cortical morphology.

## Brain imaging data

In this study, a large library of fetal and neonatal cortical surfaces reconstructed from structural T2-weighted MRI was used to derive the cortical parcellation and construct subject-specific computational brain models. We used the imaging data from the fourth data release of the developing Human Connectome Project (dHCP) (https://www.developingconnectome.org). Fetal MRI was acquired on a 3-T Philips Achieva scanner using a 32-channel cardiac coil, whereas neonatal MRI was acquired on the same scanner platform using a dedicated 32-channel neonatal head coil^52^. Following availability and quality control of the anatomical images, we retained the fetal cohort comprised 242 scans from 231 participants (123 male, 108 female), covering 21-38 weeks of postmenstrual age. For neonatal dataset, we excluded preterm-born neonatal participants, defined as those born before 37 postmenstrual weeks^53^, and retained 594 scans from 589 participants (318 male and 271 female), spanning 37-45 postmenstrual weeks. The final analytical cohort therefore comprised 836 imaging sessions from 747 unique participants, covering 21-45 postmenstrual weeks. Detailed MRI acquisition protocols and cohort-selection criteria are provided in the Supplementary Materials.

### Developmental cortical parcellation

We reconstructed cortical surfaces from the T2-weighted images using the dHCP structural processing pipeline^54^ and retained the white, pial and mid-thickness surfaces for each hemisphere. Each individual surface was registered to the corresponding age-matched dHCP cortical template using cortical folding information, establishing vertex-wise correspondence across participants and postmenstrual ages. For participants with repeated examinations, longitudinal registration was performed sequentially between adjacent time points before mapping to the age-matched template, so that later scans were aligned through the preceding earlier scan and within-subject correspondence was preserved across development. the registered cortical surfaces were further downsampled to 10,242 vertices per hemisphere before developmental cortical regionalization.

Following cortical surface reconstruction and registration, vertex-wise cortical surface area was computed on the mid-thickness surface. For vertex *i*, the local surface area was defined as

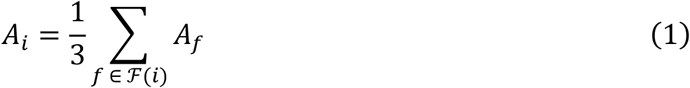

where ℱ (*i*) is the set of triangular faces incident to vertex *i*, and *A*_*f*_ is the area of face *f*, thus each triangular face contributes equally to each of its three vertices^13^. After exclusion of the medial wall, the surface-area map of each scan was normalized by its mean cortical surface area to reduce inter-individual differences in overall cortical size. The normalized surface-area maps were further assembled into a vertex-by-scan matrix after ordering scans by increasing postmenstrual age. Thus, each row, denoted ***v***_*i*_, represents the age-ordered developmental surface-area profile of cortical vertex *i*.

To characterize similarities between cortical developmental profiles, two complementary similarity matrices ***S***_1_ and ***S***_2_ were constructed:

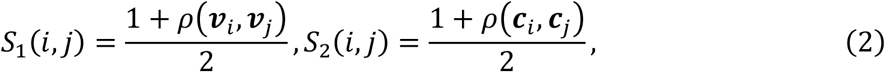

where *ρ*(∙,∙) denotes the Pearson correlation coefficient, and ***c***_*i*_ = [*ρ*(***v***_*i*_, ***v***_*r*1_), …, *ρ*(***v***_*i*_, ***v***_*rR*_)] denotes the correlation profile of vertex *i* with a set of approximately uniformly distributed reference vertices, *R* is the number of reference vertices, as selected by farthest-point sampling on the cortical surface. Thus, ***S***_1_ captures direct similarity between developmental profiles, whereas ***S***_2_ captures higher-order similarity by comparing their relationships to distributed cortical reference locations^12^. The two similarity matrices were further integrated using similarity network fusion (SNF), as seen in Fig. 1b. For each ***S***_*n*_, *n* ∈ {1,2}, a full normalized kernel ***K***_*n*_ and a sparse neighborhood kernel ***Q***_*n*_ were defined as

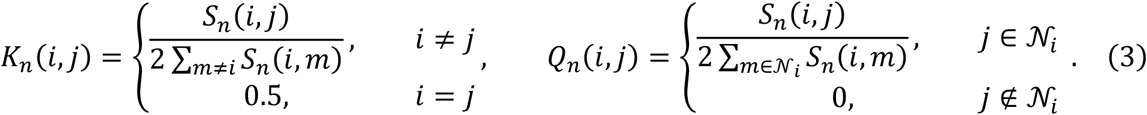

Here, *N*_*i*_ denotes the set of *M* vertices with the highest similarity to vertex *i*. Accordingly, ***K***_*n*_ preserves the global similarity structure, whereas ***Q***_*n*_ restricts information propagation to the most similar vertices. The two similarity networks were then iteratively fused using message-passing method,

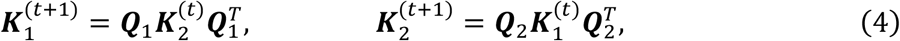

where (∙) indicates matrix transpose. This iterative propagation reinforces similarities supported by both developmental representations while attenuating weak or inconsistent relationships. After *N*_iter_ iterations, the fused similarity matrix was obtained as average of ***K***_1_ and ***K***_2_, 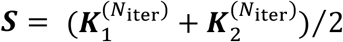. Spectral clustering was subsequently applied to the fused similarity matrix ***S*** to identify cortical regions with distinct developmental surface-area patterns. In this study, *R* = 300 reference vertices were sampled per hemisphere, the *M* = 50 most similar vertices were retained in the sparse SNF kernel, and the fusion procedure was iterated *N*_iter_ = 3 times.

Parcellation quality was evaluated using the silhouette coefficient and split-half instability. The silhouette coefficient was defined as *s*(*i*) = (*b*(*i*) − *a*(*i*))/max{*a*(*i*), *b*(*i*)} for each vertex *i*, where *a*(*i*) is the mean within-region dissimilarity and *b*(*i*) is the minimum mean dissimilarity to any other region. Dissimilarity was defined using correlation distance between developmental surface-area profiles. The mean silhouette coefficient across sampled cortical vertices was used as the overall clustering-quality measure, with larger values indicating greater within-region similarity and stronger between-region separation^13^.

Parcellation stability was assessed using repeated split-half analysis. For each candidate region number, imaging sessions were randomly divided into two subsets, the complete parcellation procedure was independently repeated for each subset, and the resulting partitions were compared with the full-data parcellation using the adjusted Rand index. This procedure was repeated across five random splits, and the mean disagreement was used as an instability measure; lower instability therefore indicates greater robustness of the parcellation to variation in the imaging sample^26^.

### Regional growth-law library

To translate population-level cortical development into growth functions for computational modeling, we constructed a regional growth law library from the developmental surface-area trajectories of the identified cortical regions, as shown in Fig. 1c. For each cortical region and hemisphere, regional surface area was modeled as a continuous function of postmenstrual age (PMA) using a generalized additive mixed model (GAMM). The model was expressed as *A*_*ij*_ = *β*_0_ + *f*(PMA_ij_) + *b*_*i*_ + *ϵ*_*ij*_, where *β*_0_ is the population intercept, *f*(PMA) is a smooth nonlinear function describing age-related changes in regional surface area, *b*_*i*_ is a subject-specific random intercept accounting for repeated measurements, and *ϵ*_*ij*_ denotes residual variation. The population-level fitted trajectory, 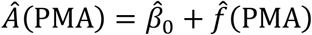, was used to characterize the typical developmental pattern of each cortical region. This framework accommodates irregular age sampling and repeated observations while capturing nonlinear developmental trends across the imaging population.

The fitted continuous GAMM trajectories allow regional growth to be derived over arbitrary developmental intervals. For a simulation spanning postmenstrual ages PMA_0_ to PMA_1_, age was rescaled to a dimensionless simulation time, *t* = (PMA − PMA_0_)/(PMA_1_ − PMA_0_), such that *t* ranges from 0 to 1 over the selected developmental window. Assuming approximately isotropic expansion within each local region of the cortical tangent plane, the regional surface-area trajectory was converted to the corresponding tangential growth ratio as 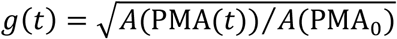.

With this normalization, *g*(0) = 1 while 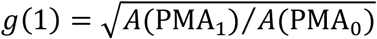 represents the accumulated tangential growth over the selected interval. Normalizing each trajectory to its window-specific starting age allows the same formulation to be applied to simulations initiated at different developmental stages. This is particularly advantageous for patient-specific modeling, where the initial postmenstrual age may vary between cases.

The GAMM-derived growth trajectories were then converted into explicit analytical expressions for computational implementation. Symbolic regression was used to approximate each normalized trajectory with a polynomial function of simulation time. Restricting the search space to polynomial expressions provided compact and continuously differentiable growth laws that could be directly implemented in the finite-element model.

The above procedure was repeated across cortical regions, hemispheres, and developmental windows to construct the regional growth law library. Because the GAMM trajectories are continuous with respect to postmenstrual age, growth laws for different simulation windows can be obtained directly from the corresponding segments of the fitted population trajectories. Each segment is then renormalized to its starting age. In this way, the library captures both regional heterogeneity and age-dependent cortical expansion. The resulting growth laws were used as regional growth controls in the subsequent whole-brain simulations. Additional details on GAMM fitting and symbolic regression are provided in the Supplementary Materials.

### Biophysical model of cortical folding

Cortical folding is shaped by coupled biological and mechanical processes. From a mechanical perspective, it can be described as a growth-induced instability driven by differential expansion between the developing cortex and the underlying subcortex. We modeled this process within the framework of finite-growth continuum mechanics, where the total deformation gradient ***F*** is multiplicatively decomposed into a growth component ***F***^*g*^ and an elastic accommodation ***F***^*e*^:

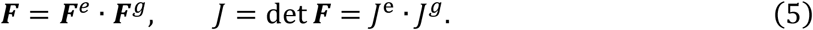

Here, *J*^*e*^ = det(***F***^*e*^) and *J*^*g*^ = det(***F***^*g*^) denote the elastic and growth-induced volume changes, respectively. Cortical growth was modeled as purely tangential expansion, while radial growth was neglected because of its comparatively limited influence on cortical folding^18, 55^. The corresponding cortical growth tensor was defined as:

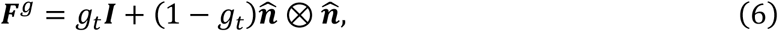

where ***I*** is the identity tensor, 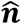 denotes the local outward surface normal, and *g*_*t*_ = *g*(*t*) is the tangential growth ratios obtained from the growth-law library described above.

The elastic response of both cortex and subcortex was described using a nearly incompressible neo-Hookean hyperelastic model. The strain-energy density was expressed in terms of the elastic deformation gradient as

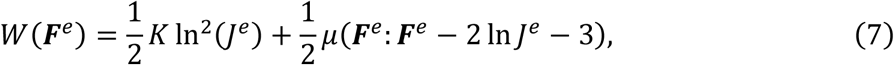

where *K* and *μ* denote the bulk and shear moduli, respectively. From thermodynamic principles, the first Piola-Kirchhoff stress ***P*** is work-conjugate to the total deformation gradient ***F***. The elastic first Piola-Kirchhoff stress is obtained from ***P***^*e*^ = ∂*W*(***F***^*e*^)/∂***F***^*e*^, and the total first Piola-Kirchhoff stress follows from the multiplicative decomposition as ***P*** = ***P***^*e*^ ∙ ***F***^*g*−*T*^. Mechanical equilibrium was governed by the balance of linear momentum. In the absence of body forces, this reduces to:

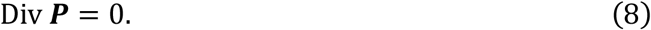

Together, differential growth between the cortex and subcortex generates incompatible local growth states that must be elastically accommodated within the connected brain tissue. The resulting stresses can drive mechanical instability and the emergence of cortical folds.

In addition to prescribed cortical growth, we incorporated stress-dependent growth in the subcortex to represent its capacity to remodel under mechanical loading. Experimental studies have shown that axons, a major constituent of developing subcortical tissue, can elongate under sustained tension and retract under compression^56^. At the continuum scale, this behavior can be represented by the stress-dependent growth law

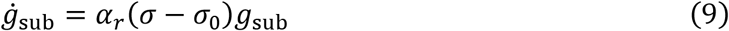

where *g*_sub_ is the subcortical growth ratio, *α*_*r*_ controls the rate of remodeling, σ is the local stress, and σ_0_ is the target or homeostatic stress. Under sustained loading, the subcortical tissue progressively remodels until the local stress approaches its target value^57^. Following the established analogy between stress-dependent growth and long-timescale viscoelastic relaxation^23^, we represented the subcortical remodeling response using a generalized Maxwell model. The time-dependent shear modulus was described using a single-term Prony-series formulation,

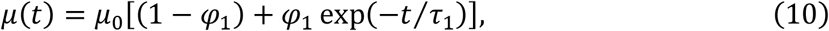

where *μ*_0_ is the instantaneous shear modulus, *φ*_1_ is the fraction of the shear response that relaxes, and *τ*_1_ is the characteristic stress relaxation time. This formulation allows stress generated by differential cortical growth to progressively relaxes within the subcortex and provides an effective continuum representation of stress-dependent growth and remodeling over developmental timescales. The cortical layer remained hyperelastic and followed the prescribed heterogeneous growth laws derived from imaging data.

### Finite-element model of the developing brain

We constructed subject-specific finite-element models from reconstructed cortical and subcortical surfaces at the initial postmenstrual age of each simulation, as seen in Fig. 1d. The framework was not restricted to a fixed starting age, for example, the cases in Fig. 4 were initialized from brain geometries spanning 21-24 weeks PMA. The cortical and subcortical surfaces were reconstructed into a conforming whole-brain solid and discretized with hexahedral elements. A nominal element size of 0.4 mm generated approximately 1.2 million elements per whole-brain model and provided at least five elements across the cortical thickness. Interior elements retained a regular voxel-like structure, whereas boundary elements were subdivided, reprojected and smoothed to preserve anatomical conformity. Cortical and subcortical elements shared nodes and faces along their interface to ensure mechanical continuity. The previously derived cortical parcellation was transferred to the finite-element model to prescribe regional growth, with 20 regions in each hemisphere. A local material orientation was defined for each cortical element to assign the tangential growth.

Brain growth and folding were simulated in ABAQUS 2023 (Dassault Systems, Paris, France) using the Dynamic Explicit solver. The cortex and subcortex were modeled as nearly incompressible neo-Hookean materials with shear moduli of 0.65 and 0.68 kPa, respectively^35^. To capture the stress-mediated growth response of the subcortex described above, we additionally represented the subcortex as a viscoelastic material using a Prony-series formulation (Eq. 10). The Prony parameter was set to *φ*_1_ = 0.9, corresponding to relaxation of 90% of the instantaneous shear response, and *τ*_1_ = 0.5. Detailed parameter settings are provided in the Supplementary Table S5. Region-specific cortical growth laws from the growth-law library were implemented through the user-defined subroutine (*VUEXPAN*), given the analogy between tissue growth and thermal expansion^58^. Cortical growth was spatially heterogeneous, with region-specific growth laws prescribed from the growth library along the local tangential directions and modulated across cortical depth using a sigmoid profile^15^, while the subcortex underwent isotropic volumetric growth (Supplementary Fig. S5). ABAQUS general contact was used for cortical self-contact, surface-to-surface contact and interactions with the boundaries. All contacts were frictionless and enforced a no-penetration condition. Simulations were performed on a Dell Alienware workstation equipped with an Intel Core Ultra 9,285K processor and 64 GB RAM. A representative simulation from approximately 21 to 43 weeks PMA required about 6 hours.

### Postprocessing and morphological quantification

Simulation outputs were analyzed using in-house postprocessing codes. At selected simulation stages, the nodal coordinates of the gray and white matter surfaces were extracted, and the corresponding cortical surfaces were reconstructed using the updated coordinates and the preserved surface connectivity. To establish correspondence with anatomical data, the undeformed simulation surface was aligned to the corresponding initial brain surface using multimodal surface matching^28^ and mapped to the standard fsLR surface coordinate system. Because the computational simulation preserved the cortical mesh topology and vertex identities, this correspondence and the associated regional labels were directly propagated to all subsequent simulation states. This procedure enabled consistent vertex-wise and regional comparisons throughout folding development and allowed cortical morphology to be reconstructed at arbitrary simulation time points.

We quantified folding morphology using complementary global, regional and landmark-based measures, including the global gyrification index (GI), sulcal depth, major sulcus count and 3-hinge gyral count. The same postprocessing procedures were applied to simulated and anatomical surfaces to enable direct comparison. Major sulci were identified using a curvature-based approach. Mean curvature was first smoothed over the cortical mesh, and spatially connected regions with curvature below −0.05 were identified as candidate sulcal basins, while small isolated components were removed using vertex-number and surface-area criteria. Notably, this analysis was designed to capture prominent sulcal structures rather than small secondary or tertiary indentations. To characterize gyral organization, we further extracted 3-hinge gyri, defined as locations where gyral crest lines converge from three directions^59^. These structures are evolutionarily preserved across primate species and exhibit distinctive structural connectivity and functional organization, making them informative landmarks for studying normal cortical architecture and its alterations in brain disorders^34^. Following established methods, the cortical surface was separated into gyral and sulcal regions based on gyral altitude, and a tree-marching procedure was used to construct crest-line trajectories within the gyral regions. Short peripheral branches were removed to obtain the principal gyral network (GyralNet), and junctions connecting three gyral branches were identified as 3-hinge gyri^59^.

For the landmark analysis in Fig. 4b, eight representative primary sulci were manually identified and visualized in ParaView 5.13.0-RC2 to illustrate the anatomical correspondence between simulated and reference brains. We additionally quantified primary sulci reproduction using the Destrieux atlas. Building on the major sulci identified above, each detected sulcal basin was assigned to the corresponding Destrieux sulcal region. A Destrieux sulcus was considered reproduced when the detected major sulcal basin covered at least 50% of the corresponding atlas-defined region. The sulci reproduction rate in Fig. 5a was quantified as (*N*_simu_/*N*_real_ * 100%), where *N*_simu_ is the number of sulci reproduced in simulation, *N*_real_ is the total number of reference sulci evaluated in matured anatomical brains. We considered 34 Destrieux sulcal regions per hemisphere^27^.

## Statistical analysis

Statistical analyses were performed to compare the numbers of major sulcal basins (Fig. 4d) and 3-hinge gyri (Fig. 4e) between anatomical and simulated brains. Left and right hemispheres were analyzed separately. Group differences were assessed using two-sided permutation tests of the difference in arithmetic means with 100,000 permutations. Raw two-sided *P* values are reported, and *P* < 0.05 was considered statistically significant. All analyses were performed using in-house Python scripts.

## Data Availability

The data supporting these findings, including the model files and regional surface area measurements used to characterize the growth model, have been deposited in the Figshare repository and will be made publicly available upon publication. The MRI data obtained from the dHCP dataset are publicly available through the Developing Human Connectome Project repository at https://www.developingconnectome.org.

## Code Availability

The code supporting this study, including scripts for growth model characterization, computational model construction, simulation postprocessing, data analysis and figure creation, is available on GitHub at https://github.com/BioDMX-UGA/Grow2Fold.

## Acknowledgements

Heterogenous growth characterization were achieved using data made available from the Developing Human Connectome Project funded by the European Research Council under the European Union’s Seventh Framework Programme (FP/2007-2013) / ERC Grant Agreement no. [319456]. We are grateful to the families who generously supported this research.

## Author Contributions Statement

JH: Methodology, Software, Validation, Investigation, Writing – Original Draft; AS: Data Curation, Methodology, Validation, Writing – Original Draft; KJ, BG, TL, XC, TW, KS, YL, WS, DZ, TL, EK, GL: Validation, Writing – Review and Editing; MJR: Conceptualization, Validation,Writing – Original Draft, Writing – Review and Editing; XW: Conceptualization, Validation, Supervision, Funding acquisition, Writing – Original Draft, Writing – Review and Editing.

